# Impaired EAT-4 Vesicular Glutamate Transporter Leads to Defective Nocifensive Response of *Caenorhabditis elegans* to Noxious Heat

**DOI:** 10.1101/620302

**Authors:** Sophie Leonelli, Bruno Nkambeu, Francis Beaudry

## Abstract

In mammals, glutamate is an important excitatory neurotransmitter. Glutamate and glutamate receptors are found in areas specifically involved in pain sensation, transmission and transduction such as peripheral nervous system, spinal cord and brain. In *C. elegans*, several studies have suggested glutamate pathways are associated with withdrawal responses to mechanical stimuli and to chemical repellents. However, few evidences demonstrate that glutamate pathways are important to mediate nocifensive response to noxious heat. The thermal avoidance behavior of *C. elegans* was studied and results illustrated that mutants of glutamate receptors (*glr-1, glr-2, nmr-1, nmr-2)* behaviors was not affected. However, results revealed that all strains of *eat-4* mutants, *C. elegans* vesicular glutamate transporters, displayed defective thermal avoidance behaviors. Due to the interplay between the glutamate and the FLP-18/FLP-21/NPR-1 pathways, we analyzed the effectors FLP-18 and FLP-21 at the protein level, we did not observebiologically significant differences compared to N2 (WT) strain (fold-change < 2) except for the IK602 strain. The data presented in this manuscript reveals that glutamate signaling pathways are essential to elicit a nocifensive response to noxious heat in *C. elegans*.

## Introduction

In mammals, glutamate is a major excitatory neurotransmitter used by primary afferent synapses and neurons located in the spinal cord [1-3]. Additionally, glutamate and glutamate receptors are found in areas specifically involved in pain sensation, transmission and transduction including the peripheral nervous system, the spinal cord and the brain [4]. Thus, glutamate receptors were investigated as a target for the development of new analgesics, but the extensive distribution and physiological functions may explain observed undesirable effects. *Caenorhabditis elegans* (*C. elegans*) consists of 959 cells including 302 neurons, making this model attractive to study neuronal communication at the physiological level [5]. *C. elegans* is specifically suitable to study nociception as it exhibits a well-defined and reproducible nocifensive behavior, involving a reversal and change in direction away from the noxious stimulus [5-7]. Consequently, *C. elegans* is a commonly used model organism to examine heat avoidance [5,7-9]. Following the genome sequencing of *C. elegans*, some genes were classified as Transient Receptor Potential (TRP) ion channel proteins with important sequence homologies to mammalian TRP channels including Transient Receptor Potential Vanilloid Channels (TRPVs) [5]. In *C. elegans*, TRPV orthologs (e.g. OSM-9 and OCR-2) were identified and it was revealed they are associated with sensory transduction [25,26]. Several studies have shown that temperature ranging from 30°C to 35°C are adequate to study the nocifensive response of *C. elegans* associated with the activation of TRPV orthologs OSM-9 and OCR-2 [5,25,27,28]. Recently, we have demonstrated *C. elegans* avoidance to noxious heat is strongly associated with the FLP-18/FLP-21/NPR-1 neuropeptide signaling pathways but results also suggested that other neuropeptides signaling pathways or classical neurotransmitters are likely playing an important role [7]. The analysis of glutamate receptor function in *C. elegans* has been previously investigated [10-12]. Neuronal communications through chemical synapses involve the activation of several neurotransmitter receptors on postsynaptic interneurons and neurons, including glutamate receptors in *C. elegans*. Studies have suggested glutamate pathways are associated with withdrawal responses to mechanical stimuli and chemical repellents [13,14]. However, the results of one study using a thermal avoidance assays suggested glutamatergic neurotransmission could be important to mediate nocifensive responses to noxious heat in *C. elegans* [5]. The glutamate needs a specific transporter to move across membranes and participate in chemical synapses. *C. elegans* vesicular glutamate transporters (i.e. *eat-4*) is therefore necessary for glutamatergic synaptic transmission [15]. Nevertheless, further validation is necessary especially since thermal avoidance behavior appears to be modulated by glutamate and neuropeptides.

## Experimental

### Chemicals, Reagents and *C. elegans* strains

All chemicals and reagents were obtained from Fisher Scientific (Fair Lawn, NJ, USA) or MilliporeSigma (St-Louis, MO, USA). For mass spectrometry analysis, formic acid, water (HPLC-MS Optima grade), acetonitrile (HPLC-MS Optima grade), trifluoroacetic acid (TFA), were purchased from Fisher Scientific. The N2 (Bristol) isolate of *C. elegans* was used as a reference strain. Mutant strains used in this work included: *grl-1* (KP4); *glr-2* (RB1808); *nmr-1* (VM487); *nmr-2* (VC2623); *flp-18* (AX1410); *flp-21* (RB982); *npr-1* (CX4148); *eat-4* (IK600); *eat-4* (IK602); *eat-4* (MT6308); *eat-4* (MT6318). N2 (Bistrol) and other strains were obtained from the *Caenorhabditis* Genetics Center (CGC), University of Minnesota (Minneapolis, MN, USA). Strains were maintained and manipulated under standard conditions as described [16,17]. Nematodes were grown and kept on Nematode Growth Medium (NGM) agar at 22°C in a Thermo Scientific Heratherm refrigerated incubators. Analyses were performed at temperature ranging from 22 to 25 °C unless otherwise noted.

### Thermal avoidance assays

The principle behind evaluating the *C. elegans* response to a stimulus (i.e thermal or chemical) is to observe and quantify the movement evoked in response to a specific stimulus. The method proposed in this manuscript for the evaluation of thermal avoidance behavior is adapted from the two and four quadrants strategies previously described [17,18]. We have previously published the experimental details for the behavioral assay used in this manuscript [7]. The technical schematics of the four quadrants assay used are described in Fig 1. Briefly, experiments were performed on 92 × 16 mm petri dishes divided into four quadrants. A delimited middle circle (i.e. 1 cm diameter) is an area where animals were not counted. The quadrants create an alternating configuration between thermal stimuli areas (i.e. 32°C) and control areas (i.e. 22°C) to prevent any bias that may occur resulting from the original location of the nematodes. Petri dishes were divided into quadrants; two stimulus areas (A and D) and two control areas (B and C). Sodium azide (i.e. 0.5M) was used in all quadrants to paralyze the nematodes but was added axially 0.5 cm behind the stimulus point to avoid paralyzing the nematode before it can display an avoidance behavior. Nociceptive heat was created using an electronically heated metal tip (0.8-mm diameter) producing a constant radial temperature gradient (e.g. 32-35°C on the NGM agar at 2 mm from the tip measured with an infrared thermometer). Nematodes manipulations were carried out according to protocol outlined by Margie *et al*. [17]. Nematodes were off food pending and throughout all experimentations. The nematodes (i.e. 50 to 200) were placed at the center of a marked petri dish and after 30 minutes, they were counted per quadrant. The derived Thermal avoidance Index (TI) formula is presented in Fig 1. Both TI and the nematode avoidance percentage were used to phenotype all tested *C. elegans* strains. The selection of quadrant temperature was based on previous experiments [5].

**Figure 1.**
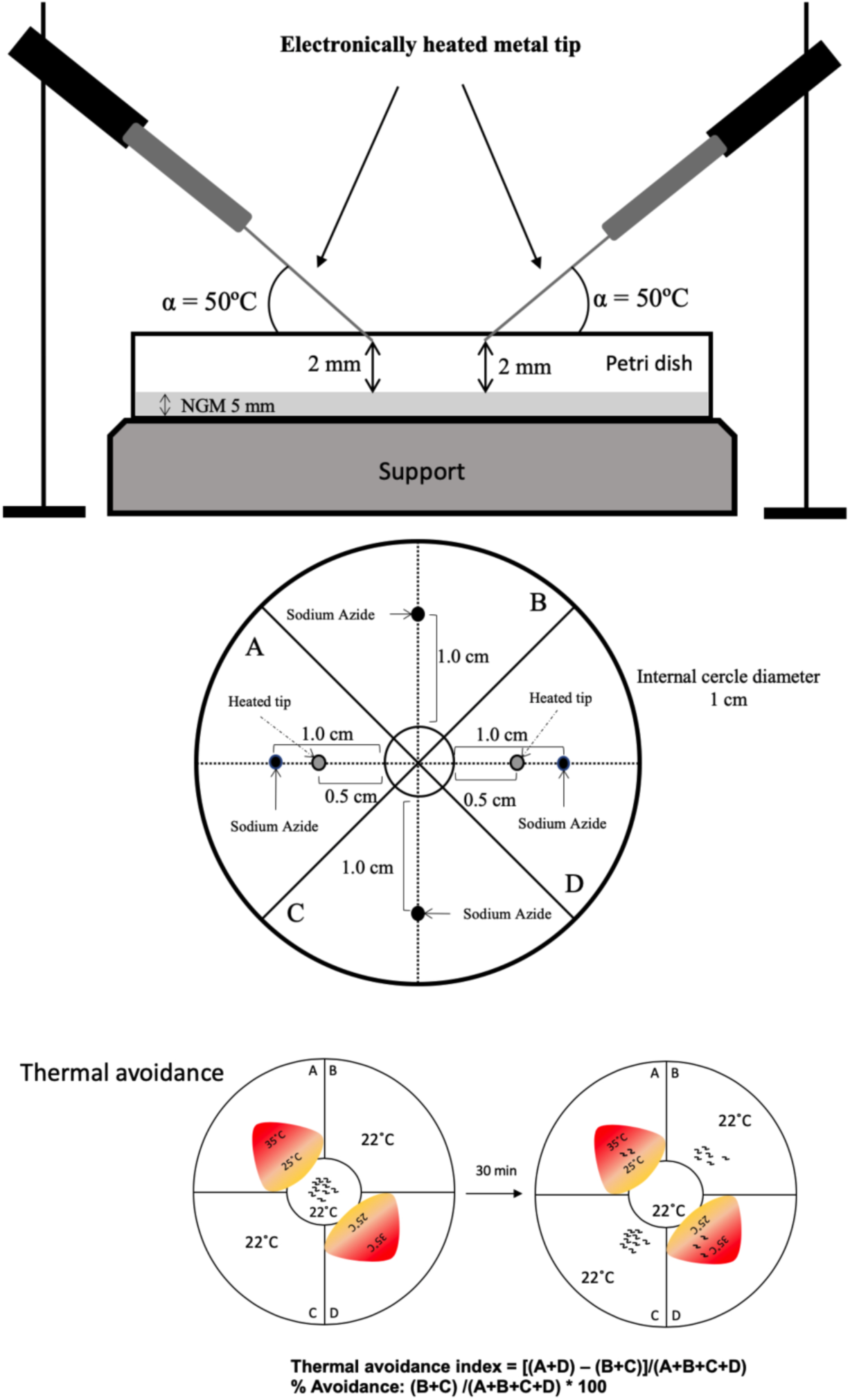
Technical schematics of the four quadrants assay adapted from Margie *et al*. (2013). For thermal avoidance assay, plates were divided into quadrants two test (A and D) and two controls (B and C). Sodium azide was added to all four quadrants to paralyse nematodes. *C*.*elegans* were added at the center of the plate (n=50 to 200) and after 30 minutes, nematodes were counted on each quadrant. Only nematodes outside the inner circle were scored. The calculation of the thermal avoidance index was performed has outlined. Quadrants temperature was continuously monitored using an infrared thermometer.

### Targeted Proteomic Analysis

Mutant strains used for the proneuropeptide analyses work included: N2 (Bristol); *eat-4* (MT6308); *eat-4* (MT6318); *eat-4* (IK600); *eat-4* (IK602). Strains were cultured in liquid media standard as described [17,18]. The liquid media was centrifuged at 1,000 *g* for 10 min and nematodes were collected and aliquoted to re-enforced 1.5 mL homogenizer tubes containing 500 µm glass beads. A solution of 8 M urea in 100 mM TRIS-HCL buffer (pH 8) containing cOmplet™ protease inhibitor cocktail (Roche) was added at a ratio of 1:5 (w:v). Lysing and homogenization was performed with a Fisher Bead Mill Homogenizer set at 5 m/s for 60 seconds and repeated 3 times with a 30 second delay. The homogenates were centrifuged at 9,000 *g* for 10 min. The protein concentration for each homogenate was determined using a Bradford assay [21]. Two hundred µg of proteins were aliquoted for each sample and proteins were extracted using ice-cold acetone precipitation (1:5, v/v). The protein pellet was dissolved in 100 µL of 50 mM TRIS-HCl buffer (pH 8) and the solution was mixed with a Disruptor Genie used at maximum speed (2,800 rpm) for 15 minutes and sonicated to improve protein dissolution yield. The proteins were denatured by heating at 120°C for 10 min using a heated reaction block. The solution was allowed to cool down 15 minutes. Proteins were reduced with 20 mM dithiothreitol (DTT) and the reaction was performed at 90 °C for 15 minutes. Then proteins were alkylated with 40 mM iodoacetamide (IAA) and the reaction was performed at room temperature for 30 min. Five µg of proteomic-grade trypsin was added and the reaction was performed at 37°C for 24h. The protein digestion was quenched by adding 10 µL of a 1% TFA solution. Samples were centrifuged at 12,000 *g* for 10 min and 100 µL of the supernatant was transferred into injection vials for analysis. The HPLC system was a Thermo Scientific Vanquish FLEX UHPLC system (San Jose, CA, USA). The chromatography was performed using gradient elution along with a microbore column Thermo Biobasic C18 100 × 1 mm, with a particle size of 5 μm. The initial mobile phase condition consisted of acetonitrile and water (both fortified with 0.1% of formic acid) at a ratio of 5:95. From 0 to 2 min, the ratio was maintained at 5:95. From 2 to 92 min, a linear gradient was applied up to a ratio of 40:60 and maintained for 3 min. The mobile phase composition ratio was reverted at the initial conditions and the column was allowed to re-equilibrate for 20 min. The flow rate was fixed at 50 µL/min and 5 µL of sample were injected. A Thermo Scientific Q Exactive Plus Orbitrap Mass Spectrometer (San Jose, CA, USA) was interfaced the UHPLC system using a pneumatic assisted heated electrospray ion source. Nitrogen was used for sheath and auxiliary gases and they were set at 10 and 5 arbitrary units. Auxiliary gas was heated to 200°C. The heated ESI probe was set to 4000 V and the ion transfer tube temperature was set to 300°C. MS detection was performed in positive ion mode and operating in TOP-10 Data Dependent Acquisition (DDA). A DDA cycle entailed one MS^1^ survey scan (*m/z* 400-1500) acquired at 70,000 resolution (FWHM) and precursors ions meeting user defined criteria for charge state (i.e. z = 2,3 or 4), monoisotopic precursor intensity (dynamic acquisition of MS^2^ based TOP-10 most intense ions with a minimum 1×10^5^ intensity threshold). Precursor ions were isolated using the quadrupole (1.5 Da isolation width) and activated by HCD (28 NCE) and fragment ions were detected in the Orbitrap at 17,500 resolution (FWHM). MS Data were analyzed using Proteome Discovered 2.2, a targeted database containing FLP-18 and FLP-21 protein sequences and a label free quantification strategy.

### Statistical Analysis

All data were analyzed using a one-way ANOVA followed by Dunnett multiple comparison test (e.g. WT(N2) was the control group used). D’Agostino-Pearson normality test was used. Significance was set a priori to p < 0.05. Additionally, only for proteomic data, the threshold is set to 2 fold-change for biological significance. The statistical analyses were performed using PRISM (version 8.1.0).

## Results and discussion

The initial experiment assessed of the mobility and quadrant bias for WT (N2) and all mutant nematodes used for this study. Nematodes were placed in the center of plates divided into quadrants conserved at constant temperature (i.e. 22°C) and no stimulus was applied (negative control). As revealed in Figure 2, there was no quadrant selection bias observed for all *C. elegans* genotypes tested. The nematodes were not preferably choosing any quadrant and were uniformly distributed after 30 minutes following the initial placement at the center of the marked petri dish. The thermal avoidance behavior of *C. elegans* was studied on petri dishes in which two opposite quadrants had a surface temperature of 32°C to 35°C and the other two were preserved at room temperature. The results illustrated in Figure 3 suggest that mutants associated with glutamate receptor *glr-1, glr-2, nmr-1, nmr-2* thermal avoidance was not affected. However, as previously demonstrated, *flp-18, flp-21* and *npr-1* mutants displayed a hampered thermal avoidance behavior [7]. *C. elegans glr-1, glr-2, nmr-1, nmr-2* single mutants were able to carry glutamatergic synaptic transmission du to functional redundancy. Moreover, these results are not surprising due to the redundant, complementary, and cooperative roles of the glutamate system and neuropeptide signaling pathways in the modulation of nociceptive responses. In mammals, glutamate and substance P, a tachykinin neuropeptide, are amongst the molecules most involved in the transmission of pain through chemical synapse occurring into the dorsal horn of the spinal cord [22,23]. As it was previously established, heat avoidance relies partly on functional NPR-1 receptors located in the RMG interneuron and both, FLP-18 and FLP-21 mature neuropeptides are ligands of NPR-1 [19]. The results showed in Figure 4 suggest that all strains of *eat-4* mutant tested displayed defective thermal avoidance behavior. These mutants have hampered glutamatergic synaptic transmission since glutamate can’t move across membranes and participate in chemical synapses. The results obtained are coherent with our initial hypothesis and complement previously reported findings using other behavior assays [5,20] revealing an important role of glutamate pathway to trigger a nocifensive response to noxious heat. We wanted to verify if these results were also an outcome of a differential expression of the FLP-18/FLP-21/NPR-1 pathways. An upregulation of FLP-18 or FLP-21 would trigger more activation of the NPR-1 receptor located on the RMG interneurons and contribute to increase thermal avoidance response. We therefore analyzed the effectors FLP-18 and FLP-21 at the protein level and as shown in Figure 5, except for strain IK602, we have not observed biologically significant differences compared to N2 (WT) strain (fold-change < 2). Interestingly, the upregulation of FLP-18 in strain IK602 did not compensate for the absence of glutamate transporter *eat-4* and shown impede nocifensive behavior to noxious heat as shown in Figure 4. Collectively, these results reveal that neuropeptide signaling FLP-18/FLP-21/NPR-1 pathways and glutamate signaling pathways are essential to elicit a nocifensive response to noxious heat in *C. elegans*. Interestingly, the role of glutamate and neuropeptide signaling pathways in the modulation of nociceptive responses is very well recognized in mammals [22,23]. Thus, we believed *C. elegans* thermal avoidance can be used as a model to study nociception and test compound libraries targeting TRPV channels in early stage of drug discovery.

**Figure 2.**
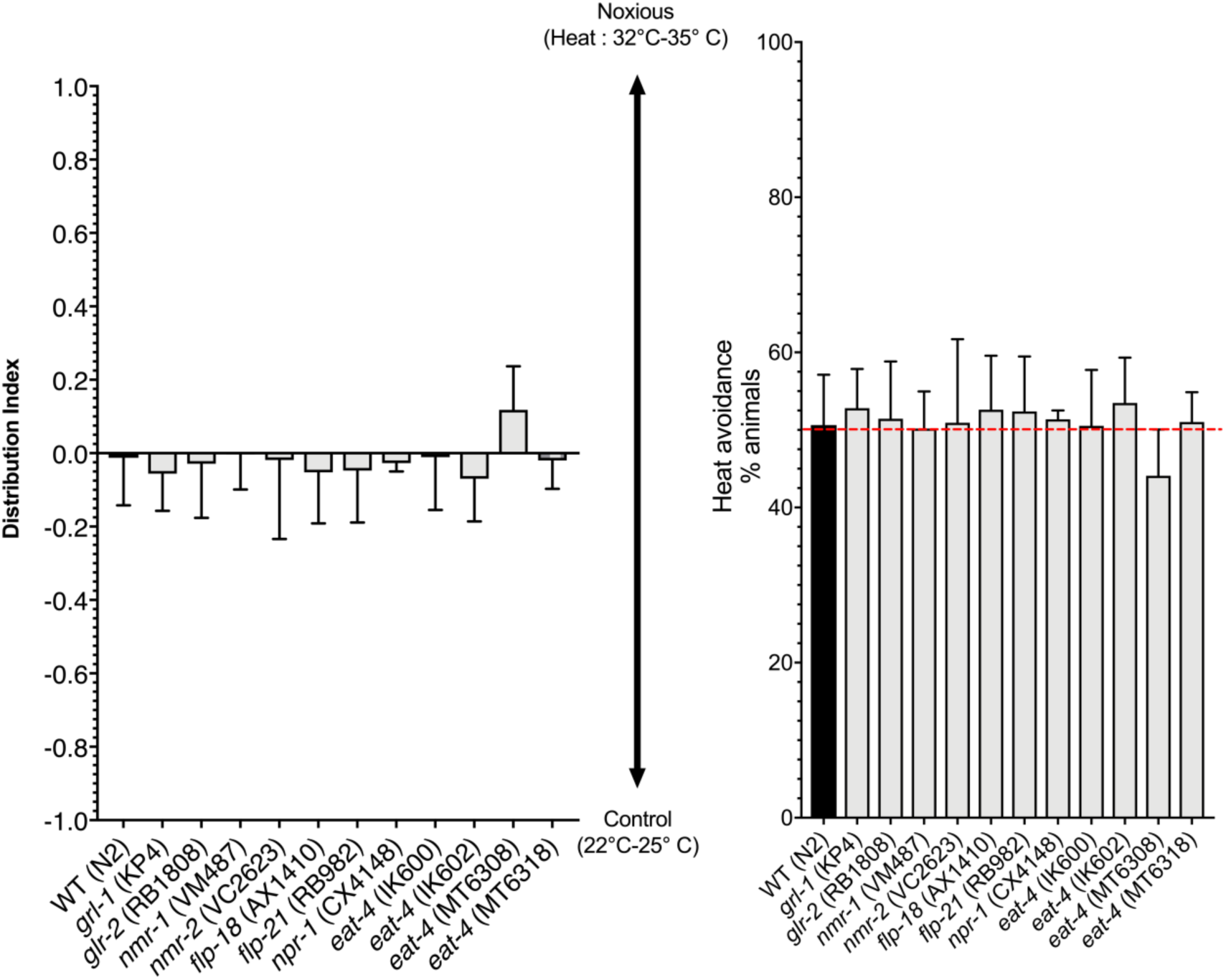
Comparison of the mobility and bias for WT (N2) and mutant *glr-1, glr-2, nmr-1, nmr-2, flp-18, flp-21, npr-1* and *eat-4* nematodes in plates divided into quadrants conserved at constant temperature and no stimulus were applied (negative control). Display values (mean ± SD) were calculated from at least 6 independent experiments (n > 300 nematodes) for each strain. No quadrant selection bias was observed for all *C. elegans* genotypes tested.

**Figure 3.**
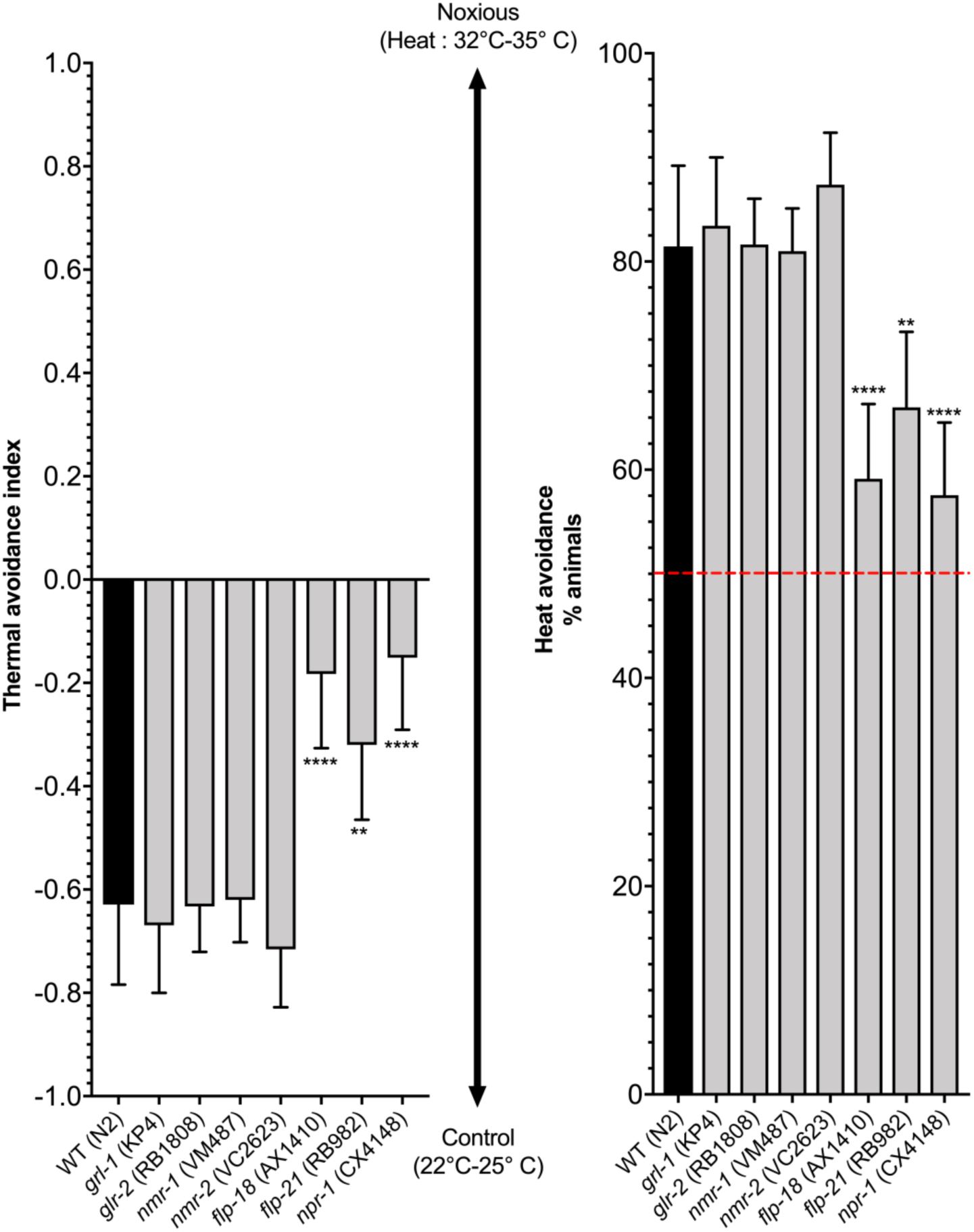
Thermotaxis index and avoidance were evaluated for WT (N2) and mutant nematodes. For thermal avoidance assay, plates were divided into quadrants two test (A and D) and two controls (B and C). Sodium azide was added to all four quadrants to paralyze nematodes. *C. elegans* were added at the center of the plate (n = 50 to 200) and after 30 min, animals were counted on each quadrant. Only animals outside the inner circle were scored. The calculation of the thermal avoidance index and chemotaxis index were performed has outlined. Quadrants temperature was continuously monitored using an infrared thermometer. Display values (mean ± SD) were calculated from at least 6 independent experiments (n > 300 nematodes) for each genotype. Heat avoidance is not affected in *glr-1, glr-2, nmr-1* and *nmr-2* but hampered in *flp-18, flp-21* and *npr-1* mutant nematodes. ** p < 0.01; **** p < 0.0001

**Figure 4.**
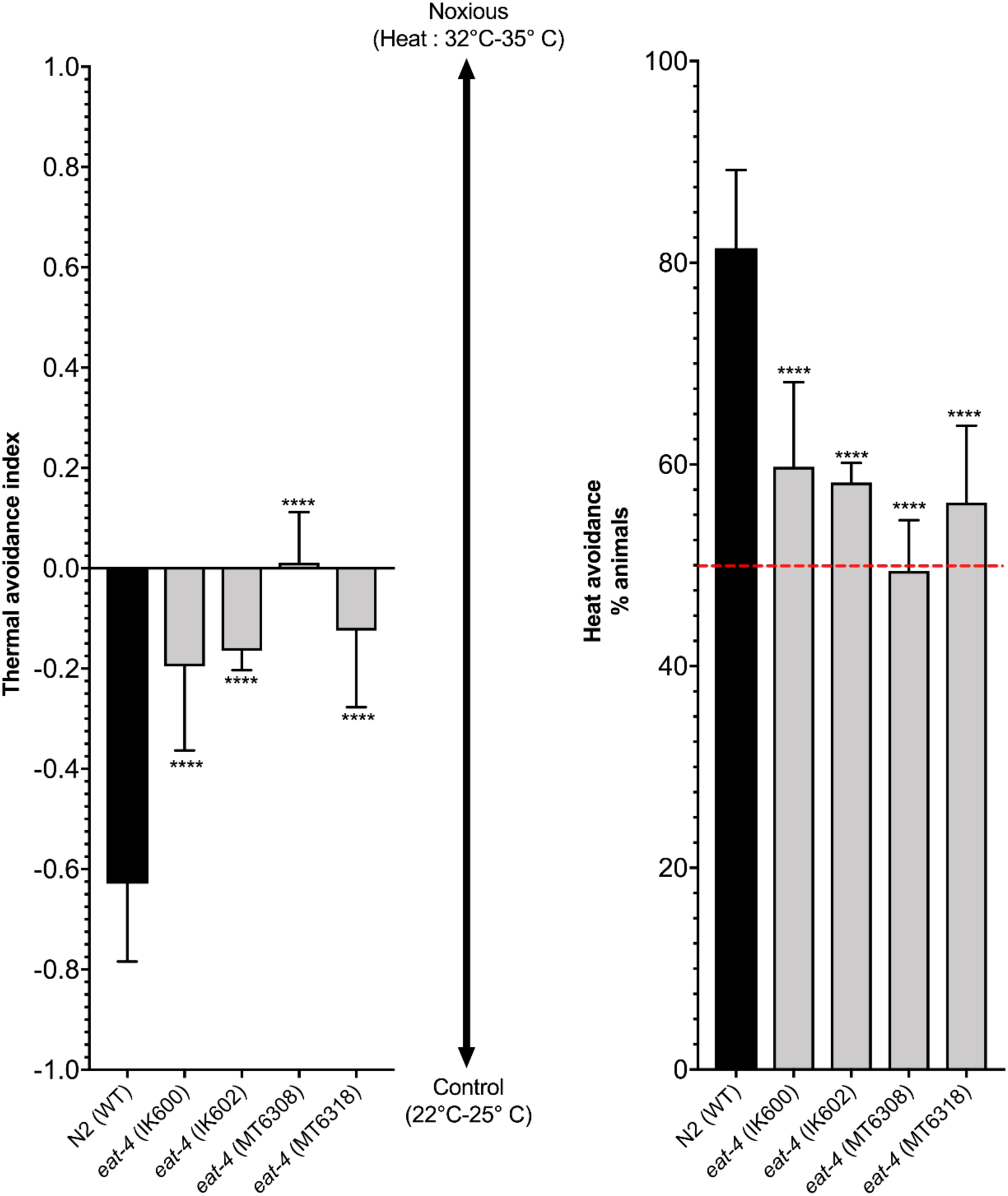
Thermotaxis index and avoidance were evaluated for WT (N2) and Vesicular glutamate transporter *eat-4* mutant nematodes. Display values (mean ± SD) were calculated from at least 6 independent experiments (n > 300 nematodes) for each genotype. Heat avoidance is impaired for *eat-4* mutant nematodes. **** p < 0.0001

**Figure 5.**
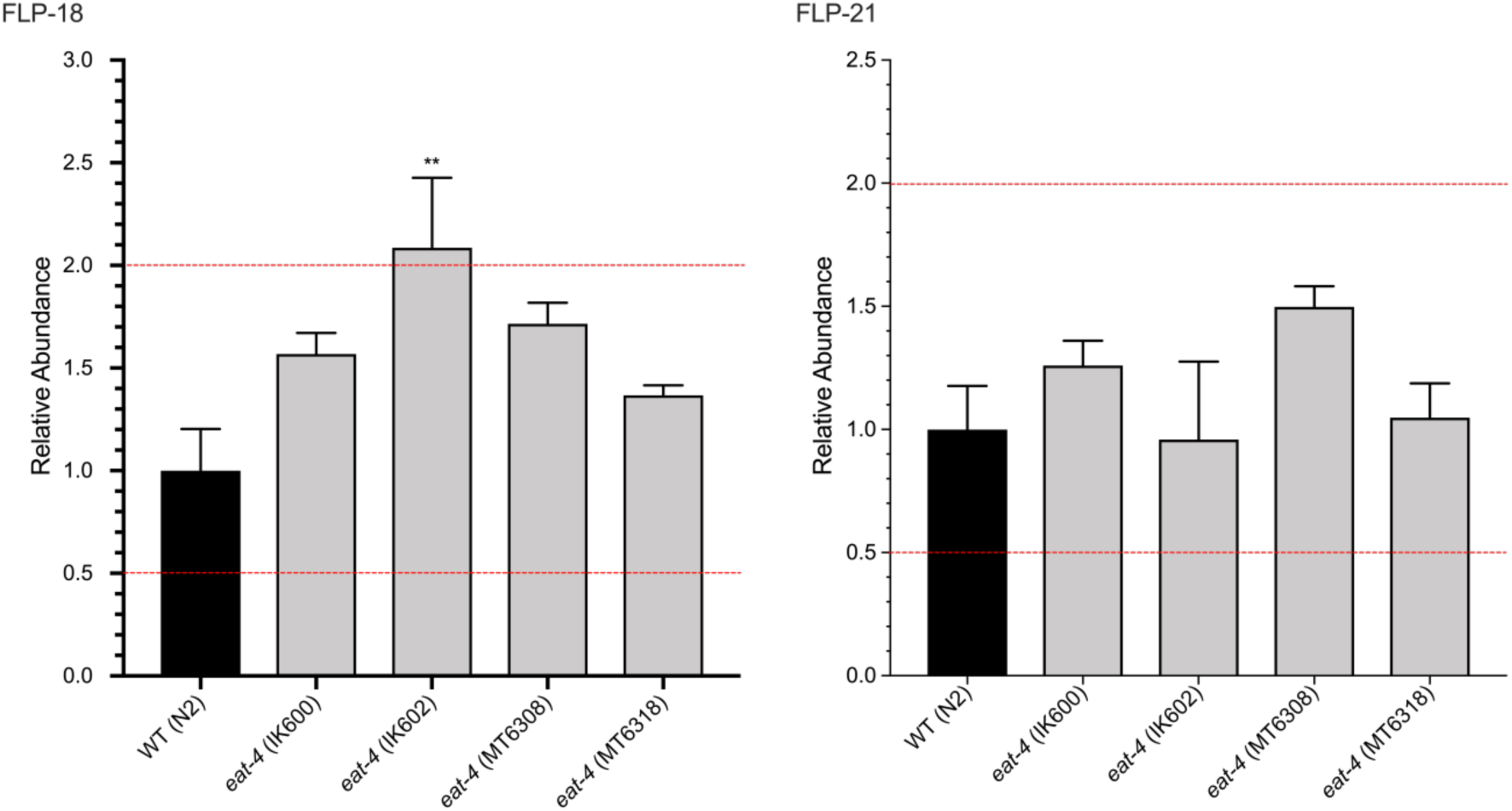
Relative concentration of FLP-18 and FLP-21 FMRF-Like peptides. The concentration of FLP-18 or FLP-21 is not significantly different (fold change < 2) for three *eat-4* mutant strains. FLP-18 is marginally different for the strain IK602. Three biological replicates were analyzed for each *C. elegans* strain and display values are mean abundance ± SD. ** p < 0.01

## Acknowledgements

This project was funded by the National Sciences and Engineering Research Council of Canada (F. Beaudry discovery grant no. RGPIN-2015-05071). Laboratory instruments were funded by the Canadian Foundation for Innovation (CFI) and the *Fonds de Recherche du Québec (FRQ)*, Government of Quebec (F. Beaudry CFI John R. Evans Leaders grant no. 36706). Sophie Leonelli received a NSERC Undergraduate Student Research Awards scholarship.

